# Healthy and diseased placenta barrier on-a-chip models suitable for high-throughput studies

**DOI:** 10.1101/2022.06.30.498115

**Authors:** Gwenaёlle Rabussier, Ivan Bünter, Josse Bouwhuis, Camilla Soragni, Chee Ping Ng, Karel Dormansky, Leon J. de Windt, Paul Vulto, Colin E. Murdoch, Kristin M. Bircsak, Henriёtte L. Lanz

**Affiliations:** MIMETAS BV, Leiden, The Netherlands; Department of Cardiology, Maastricht University, Maastricht, The Netherlands; Systems Medicine, School of Medicine, University of Dundee, Dundee, Scotland, UK

## Abstract

Preeclampsia is one of the leading causes of maternal and perinatal morbidity in the world. Its fundamental mechanisms are yet poorly understood due to a lack of good experimental models. Here we report an *in vitro* model of the placental barrier, based on co-culture of trophoblasts and endothelial cells against a collagen extracellular matrix in a microfluidic platform. The model yielded a functional syncytium with barrier properties, polarization, secretion of relevant extracellular membrane components, thinning of the materno-fetal space, hormone secretion, and transporter function. The model was exposed to low oxygen conditions and perfusion flow was modulated to induce preeclamptic characteristics, resulting in reduced barrier function, hormone secretion, and microvilli as well as increased nuclei count. The model was implemented in a titer plate microfluidic platform fully amenable to high-throughput screening. We thus believe this model could aid mechanistic understanding of preeclampsia as well as support future development of effective therapies through target and compound screening campaigns.

## Introduction

In recent years, unpredictable and untreatable pregnancy disorders have emerged as major maternal and fetal health concerns. Among those placental dysfunctions, preeclampsia (PE) is one of the leading causes of maternal and perinatal mortality and morbidity in the world^1^. It affects up to 8% of all pregnancies and contributes worldwide to 15% of maternal deaths annually^1,2^. Medical care costs associated with PE amount to an estimated $2.18 billion in the United States for the year of delivery and increase even further when long-term effects are considered^3^. PE is clinically defined by hypertension and proteinuria starting in the 20^th^ week of pregnancy. Some of the severe complications include pre-term delivery, organ damage, such as liver rupture or acute kidney failure, and cardiovascular disease^4^. Despite a rapid increase in PE incidence, as shown by a 25% increase between 1987 and 2004 in the United States^3^, there are to date few reliable and early predicative biomarkers, preventive measures, or treatment strategies available. Today no cure is available, other than premature delivery of the fetus, which can be contributed to a poor understanding of the pathogenesis of PE^5^.

It is well established that PE originates from the placenta^6^. As a unique and transient organ supporting pregnancy development, the placenta’s main role is to provide essential nutrients and gases, to mediate metabolic exchanges, as well as to protect the fetus from xenobiotics and pathogens^7^. The placenta bridges maternal and fetal circulation and is made up of a number of anatomical barriers, such as the syncytium, connective tissue and fetal capillaries^8^. The syncytium is in direct contact with maternal blood and constitutes the most important membrane barrier. It is comprised of a multinucleated layer of syncytiotrophoblasts originating from the fusion of underlying trophoblasts. The placenta is also a major source of hormone secretion, thereby playing a crucial role in maintaining a healthy 9 pregnancy^9^.

During pregnancy, the remodeling of the maternal spiral arteries by the end of the first trimester is essential for promoting adequate blood supply to the placenta and the fetus. It includes events such as trophoblast proliferation, differentiation and invasion^10^. Alterations in those events are the main hallmark of early-onset PE. Incomplete transformation of the spiral arteries leads to a reduction of the uteroplacental perfusion and subsequent placental ischemia and hypoxia^11,12^. Even though the trigger for abnormal placental development and the chronological order of events remains unknown, placental hypoperfusion is known to be accompanied by syncytium damage, impaired balance of angiogenic and anti-angiogenic factors, and release of pro-inflammatory molecules into the maternal circulation^13^.

The poor understanding of PE, lack of treatments and lack of predictive biomarkers are mainly attributed to difficulties in studying the placenta, especially at the early stage of pregnancy. *Ex vivo* placenta perfusion systems are the gold standard and have provided many clues regarding disease pathology^14^. However, they come with numerous limitations. The study of early-onset PE is hampered by restrictions in access to placenta from terminated or unsuccessful pregnancies^15^. Animal models, such as mice and sheep have been useful in modeling localized physiological processes, but none of them exactly mirrors the human placenta pathology, resulting in poor translational relevance^16,17^. *In vitro* models have been developed utilizing immortalized cell lines, primary isolated trophoblast and placenta explants^18^. Their combination with manipulation of gene and protein expression as well as oxygen tension significantly helped elucidate key features of PE^19–21^. Nevertheless, trophoblast monolayer cell culture, conventional or in Transwell systems, lacks the multilayer complexity and dynamic environment of the placenta.

Culture of placenta cells in microfluidics system brought placenta barrier modeling to a more advanced level. These organ-on-a-chip systems are enhancing tissue engineering by building more comprehensive tissues constituted of multiple cell types, extracellular matrix scaffolds, fluid flow and mechanical strain^22,23^. In the recent years, several research groups have successfully developed placenta-on-a-chip models mimicking the multi-layered maternal-fetal interface. Models were utilized to reconstitute placenta physiology and study transplacental transfer of substances such as glucose^24^, xenobiotics^25,26^, caffeine^27^ or nanoparticules^28^. Others have modeled placental inflammatory responses to bacterial infections^29^.

Current placenta-on-a-chip models do have some limitations. Several of them do not involve trophoblast cell line differentiation into syncytium^27,29–33^, which constitutes the main functional barrier of the placenta. Moreover, chips are composed of polydimethylsiloxane (PDMS), known to absorb small hydrophobic molecules such as drugs.^34^ Most importantly, these systems are typically low in throughput thus restricting the number of drugs that can be investigated. In addition, the use of microfluidics for the modeling of pregnancy disorders such as PE at the maternal-fetal interface level, has not been established yet.

Here, we report a novel, robust and high-throughput approach for modeling the human placenta barrier. The model comprises a multilayered functional syncytium and umbilical endothelium separated by a thin connective tissue-like extracellular matrix (ECM). We showed transporter functionality, barrier function and hormone secretion. To induce PE-like characteristics, we exposed our model to a hypoxic and ischemic environment. Under these conditions, the placenta barrier on-a-chip showed a damaged syncytium, characterized by reduced barrier function and differentiation, down-regulation of glucose transporter-1 expression and reduction in angiogenic factors, which are all known to be features associated with the PE pathology^6,13,35,36^. Thus, our placenta barrier on-a-chip model constitutes a promising tool for the studies of healthy and diseased pregnancy and opens the door for the evaluation of placental drug transfer and development of new therapeutics for the treatment of pregnancy disorders.

## Materials and methods

### Cell culture

Human choriocarcinoma cell line BeWo b30 clone (#C0030002; AddexBio) was cultured in T75 flasks in Optimized-DMEM (#C0003-02; AddexBio) supplemented with 10% fetal bovine serum (FBS; F4135; Sigma) and 1% penicillin-streptomycin (#15070063; Thermo Fisher). Cells were cultured in a humidified incubator (37°C; 5% CO2) and cell culture medium was changed every 2-3 days. After reaching 90% confluency, cells were trypsinized with 0.25% Trypsin-0.53mM EDTA (#30-2101, ATCC) for passaging or seeding in the OrganoPlate.

Human umbilical vein endothelial cells from pooled donors (HUVEC; #C2519AS; Lonza) were thawed from liquid nitrogen and resuspended in Complete Human Endothelial Cell medium (#H116; Cell Biologics) for seeding in the OrganoPlate.

All experiments were performed with BeWo b30 and HUVEC cells between passages 26 to 30, and passage 4 to 5, respectively. The cells were routinely tested for mycoplasma contamination and found negative.

### OrganoPlate culture

To recreate the multilayered architecture of the placenta barrier, we used the Mimetas OrganoPlate 3-lane. This microfluidic device based on a standard 384-well microtiter plate (4004-400B; MIMETAS BV) contains 40 independent chips, enabling the parallel culture of as many organ models in high-throughput (Figure 1A). Each tissue culture chip consists of three microfluidic lanes. The top and bottom perfusion lanes of 300 μm × 220 μm (w × h), and the middle lane of 350 μm × 220 μm (w × h) are all accessible by their well openings. These channels are separated on their bottom by a meniscus pinning barrier, the PhaseGuide™^37^, allowing the patterning of extracellular matrix (ECM) gel in the middle channel and the culture of cells in the bordering perfusion channels in a membrane-free manner (Figure 1B).

**Figure 1.**
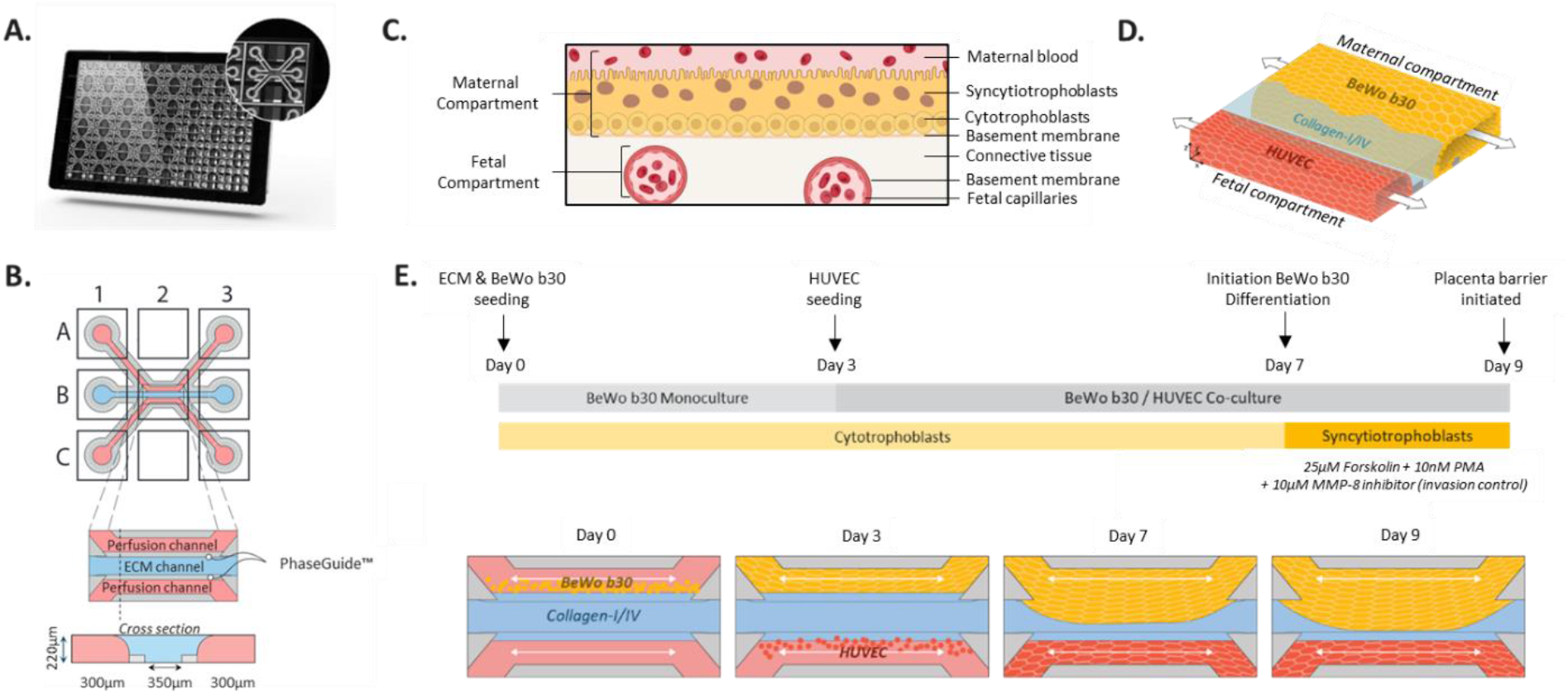
Placenta barrier on-a-chip set-up in the OrganoPlate 3-lane. (**A**) Bottom view of the OrganoPlate 3-lane, a 384-well plate format containing in its bottom 40 individual microfluidics chips. Zoomed-in image of one microfluidic chip. (**B**) Schematic representation of one chip consisting of three adjacent channels accessible by top (A1), bottom (C1) inlets and top (A3), bottom (C3) outlets wells. Perfusion channels are separated by small ridges (PhaseGuide™) which enabled the patterning of ECM gel in the middle lane and culture of cells in the perfusion channels in a membrane-free manner. Cultures are observed from the observation window (B2). (**C**) Schematic of the placenta barrier multilayered architecture. The interface is composed of syncytiotrophoblasts and cytotrophoblasts (maternal compartment) on top of their basement membrane. A thin connective tissue separates the maternal interface from the fetal capillaries (fetal compartment). (**D**) 3-dimensional schematic of the placenta barrier modeling in the OrganoPlate 3-lane, representing the maternal and fetal compartment in close proximity. Continuous flow through the maternal and fetal compartments is depicted by the arrows. (**E**) Schematic workflow depicting placenta barrier modeling set-up, BeWo b30 differentiation and controlled invasion over time.

The first-trimester placenta is characterized by a trophoblast syncytium and its underlying cytotrophoblasts separated from the fetal endothelium by a thin connective tissue^7^ (Figure 1C). To model this maternal-fetal interface, the first-trimester trophoblast human choriocarcinoma cell line BeWo b30 and HUVEC are used to recreate the maternal and fetal compartment, respectively (Figure 1D). Cells were cultured against a 3-dimensional ECM scaffold based of collagen-I and collagen-IV, key components of the placenta barrier^38^. ECM was prepared by mixing independently and respectively, in 8:1:1 ratio, rat collagen-I 8.91 mg/mL (#354249; Corning) and collagen-IV from human placenta 5 mg/mL (#234154; Sigma-Aldrich) with HEPES (1M; #15630; Life Technologies) and NaHCO_3_ (37g/L, S5761; Sigma-Aldrich) on ice. Collagen-I and collagen-IV ECM gels were then mixed in a 3:1 ratio. The collagen-I/collagen-IV ECM mixture was kept on ice and used for seeding within 10 min. Before ECM seeding, 50 μL HBSS (#H6648; Sigma-Aldrich) was added to the observation windows to prevent gel dehydration and to increase optical clarity. The ECM mixture (1.7 μL) was dispensed in the gel inlet. The plate was incubated in a humidified incubator (37°C, 5% CO_2_) for 15 min to allow the ECM to polymerize. For maternal compartment modeling, BeWo b30 cells were resuspended in their culture medium to a cell density of 12.000 cells/μL and 2 μL of cell suspension was added into the top perfusion inlet (Figure 1E). Subsequently, 50 μL of BeWo b30 culture medium was added in the top and bottom inlet wells and the plate was placed on its side for 3h to allow the cells to settle against the ECM. This was followed by addition of 50 μL medium to the top outlet, bottom inlet, and outlet wells. Bidirectional perfusion was started by placing the plate on a rocking platform (OrganoFlow; MIMETAS BV) set at a 7° inclination and 8-minute interval, in the incubator (37°C, 5% CO_2_). Medium was refreshed in top and bottom channels 2 days later. On day 3, fetal compartment modeling was initiated by HUVEC seeding into the bottom channel. Seeding of cells in channel previously wetted by medium was performed by passive pumping as previously described^39^. In short, medium was aspirated from top and bottom perfusion wells and bottom inlet to be seeded was washed with 50 μL PBS (#70013016; Thermo Fisher). 50 μL of endothelium was then dispensed on the bottom outlet and 2uL of HUVEC suspension (10.000 cells/μL) was then seeded into the previously washed inlet. To avoid dehydration, 50 μL of BeWo culture medium was dispensed in the top channel outlet well and the plate was placed on its side for 1.5h to allow the HUVEC to attach against the ECM. BeWo and HUVEC culture medium (50 μL) was respectively dispensed in top and bottom channel inlet wells. The OrganoPlate was placed back under perfusion in the incubator (37°C, 5% CO_2_). Cells were grown for several days under continuous perfusion. On day 6, medium was refreshed in top and bottom perfusion channel wells. After 7 days of culture, and to promote the differentiation of BeWo b30 into multinucleated syncytia, trophoblast cells were exposed for 48h to 25 μM Forskolin (#F3917; Sigma-Aldrich), and 10 nM of Phorbol 12-myristate 13-acetate (PMA, #P1585; Sigma-Aldrich), well known activators of trophoblast differentiation pathways^40,41^. Additionally, BeWo b30 are known to secrete a wide range of metalloproteinases, characteristic of their invasiveness and ECM remodeling^42^. Because collagen-I is the main component of the ECM gel, BeWo b30 ECM invasion behavior was controlled by supplementing trophoblast differentiation medium with 10 μM metalloproteinase-8 (MMP-8) inhibitor^43–45^. 50μL of BeWo b30 differentiation medium and HUVEC culture medium was refreshed on both days 7 and 8 in the top perfusion channel and bottom perfusion wells, respectively. Placenta barrier co-culture were fully established after 9 days of culture.

Preeclamptic characteristics were modelled by exposing placenta barrier co-cultures to a hypoxic or hypoxic-ischemic environment. Co-cultures were established as described above and moved on day 8 to a low-oxygen incubator for 24h (1% O_2_, 37°C, 5% CO_2_). To model the “Hypoxic” condition, OrganoPlate were culture on the OrganoFlow (7°, 8-min intervals), while “Ischemic” condition was mimicked by placing plates flat and static.

Phase contrast images were taken every 1 to 3 days with a high content microscope (ImageXPress XLS-C; Molecular Devices) to monitor culture development.

### Barrier integrity assay (BI Assay)

From day 7 to day 9, the tightness of the trophoblast layer was assessed daily by monitoring the flux of fluorescent dyes from the apical to basal side of the tubule. Assay was previously described in intestinal epithelium tubes^46^. To ensure proper flow profiling, the gel inlet and outlet wells were washed with 25 μL endothelium culture medium for 5 min, under perfusion. Then, medium was aspirated from all wells of the chip and 20 μL of endothelium culture medium was dispensed in middle and bottom channel wells. Subsequently, 40 μL and 30 μL of medium containing 4.4kDa TRITC-dextran (0.5mg/mL; #T1037; Sigma-Aldrich) and sodium fluorescein (10 μg/mL; #F6377; Sigma) was dispensed in the top perfusion inlet and outlet wells, respectively. The passage of dextran into the basolateral side of the trophoblast layer was monitored every 2 minutes for 12 minutes with a high content microscope (ImageXPress XLS-C; Molecular Devices) in the TRITC and FITC channels. The ratio between the fluorescent signal in the basal and apical region of the tubule (leakage score) was calculated for each timepoint using Fiji Software^47^.

### TEER measurement

Transepithelial electrical resistance (TEER) of the trophoblast barrier was measured from day 6 to day 9 using an automated multichannel impedance spectrometer (OrganoTEER, MIMETAS) as previously published, providing electrical connections to all inlet and outlets well of the OrganoPlate^48^. Before measurement, 50 μL endothelium complete medium was refreshed in all perfusion channel wells and plate was kept static at room temperature (≈ 21°C) for 30 min to ensure medium equilibration.

Using the OrganoTEER software (MIMETAS), impedance spectra of our tubes was measured (medium settings, 40 frequency points from 10Hz to 150kHz), from which the electrical resistance (Ohm) of the tubules were automatically extracted by the software fitting algorithm.

### Immunofluorescent staining

Placenta barrier cultures were fixed and prepared for immunostaining as previously described^46^. Briefly, 3.7% formaldehyde (#252549; Sigma) in HBSS (with Ca^+2^/Mg^+^) was added to the cultures for 15 min and rinsed with PBS. Cells were washed with a washing solution of 4% HI-FBS (#F4135; Sigma) in PBS and permeabilized with 0.3% Triton X-100 (#T8787; Sigma) for 10 min. Next, tubules were rinsed with washing solution and blocked with 2% HI-FBS, 2% BSA (#A2153; Sigma) and 0.1% Tween20 (#P9416; Sigma) in PBS for 45 min. The cells were then incubated with primary antibodies diluted in blocking solution overnight, static, at 4°C. The following primary antibodies were used: Mouse anti-Ezrin (#610602; BD Transduction Laboratories; 1:100), Mouse anti-ZO-1 (#33-9100; Life Technologies; 1:125), Rabbit anti-ECAD (#3195S; Cell Signaling Technology; 1:200), Mouse anti-hCG-β (#ab958; Abcam; 1:100), Mouse anti-GLUT-1 (#ab40084; Abcam; 1:125), Rabbit anti-collagen-IV (#PA5-104508; Thermo Fisher; 1:100), Rabbit anti-laminin (#ab11575; Abcam; 1:200), Rat anti-perlecan (#MA106821; Thermo Fisher; 1:100), Rabbit anti-nidogen-2 (#PA5626615, Thermo Fisher, 1:500), Mouse anti-tenascin-C (#ab88280, Abcam, 1:500), Mouse anti-fibronectin Alexa Fluor 647 Conjugated (#56098; BD Biosciences; 1:200) and Mouse isotype (#086599; Thermo Fisher). Cultures were washed three times with washing solution and then incubated for 30 min at room temperature on the OrganoFlow (7°, 2-min intervals) with Alexa Fluor 488 (#a32731; Invitrogen; 1:250) and Alexa Fluor 647 (#a31571; Life Technologies; 1:250) diluted in blocking buffer. Subsequently, cells were washed three times and nuclei were stained with Hoechst 33342 (#H3570; Thermo Fisher; 1:2000). Actin was stained using ActinGreen™ 488 ReadyProbes™ Reagent (#R37110; Thermo Fisher; 2 drops/mL) or ActinRed™ 555 ReadyProbes™ Reagent (#R37112; Thermo Fisher; 2 drops/mL) for 30 min at room temperature on the OrganoFlow (7°, 2-min intervals). Chips were imaged on the High-Content Confocal Imaging System (ImageXPress^®^ Micro Confocal; Molecular Devices). Immunostainings were processed using Fiji Software.

### Maternal-fetal distance quantification

Placenta barrier co-culture after 9 days of culture were fixed and permeabilized as previously described. Cultures were stained with ActinRed™ 555 ReadyProbes™ for 30 min on the OrganoFlow (7°, 2-min intervals). Z-stack from the bottom to the top of the chips were imaged on the High-Content Confocal Imaging System with 10x objective. Maximum fluorescence intensity projection (max projection) was performed and distance between the maternal epithelium and fetal endothelium was quantified using Fiji Software. For every chip, distances were measured in 5 locations, including the shortest and longest points between the maternal and fetal tubules and 3 fixed common points across all chips. The 5 measured distances were then averaged.

### RNA extraction and quantitative real-time PCR (RT-qPCR)

Total RNA was extracted and isolated from BeWo b30 tubules using a RNeasy Mini Kit (#74004; QIAGEN). To specifically isolate RNA from trophoblast cells, endothelium was first lysed using RLT buffer, and cell lysate was aspirated. Next, BeWo b30 from 5 individual chips were lysed, pooled, and total RNA concentration was quantified with the NanoDrop OneC^®^ spectrophotometer (Thermo Fisher). cDNA was synthetized using M-MLV reverse transcriptase (#28025013; Invitrogen). Quantitative real-time PCR (qPCR) was then performed using SYBR Green PCR Master Mix (#4309155; Thermo Fisher) with the LightCycler^®^ 480 device (Roche). PCR primer sequences used for RT-qPCR are listed in Table 1. Target gene expression levels were normalized to GAPDH or Actin. Data were analyzed with the Roche, LightCycler software version 1.1.

**Table 1.**
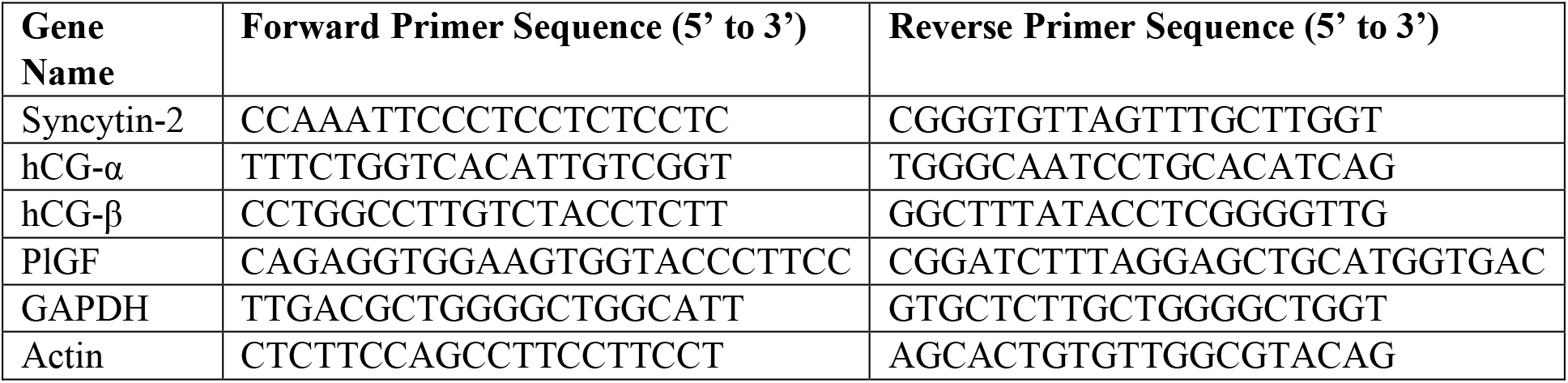
qPCR primer sequences

### Human chorionic gonadotropin Beta (hCG-β) quantification

BeWo culture medium from the top perfusion inlet and outlet wells was collected on day 7 (prior to trophoblast differentiation), day 8 (24h post-differentiation) and day 9 (48h post-differentiation) and stored at −80C° until assay was performed. The concentration of hCG-β in the culture medium was measured using the Human CG beta (HCG beta) Duoset ELISA (#DY9034-05, R&D Systems) according to the manufacturer protocol.

### Drug transport activity assays

The transport activity of multidrug resistance-associated protein (MRP) transporters was assessed in the placenta barrier on-a-chip as previously described^49^. Briefly, 30 μL of 1.25 μM 5-chloromethylfluorescein-diacetate (CMFDA, #C7025, Life Technologies) diluted in Opti-HBSS (1:2 Opti-MEM (#11058021, Gibco)) and HBSS) was added to all the perfusion inlets and outlets, either with two MRP inhibitors, MK571 (5 μM, 10 μM, 30 μM, #M7571, Sigma) and Quercetin^50,51^ (30 μM, #Q4951, Sigma)) or their vehicle (DMSO, #D8418, Sigma). OrganoPlates were incubated in a humidified incubator (37°C, 5% CO2) for 30min on the OrganoFlow (7°, 8-min intervals). Once inside the cell, CMFDA is metabolized to fluorescent compound MF-SG which is a substrate for MRPs. Cultures were then cooled down by adding 50 μL cold HBSS in the observation window. Assay solution was replaced with ice-cold inhibitor cocktail targeting MRP, BCRP and Pgp transporters (10 μM MK571, 10 μM Ko143 (#K2144, Sigma), 10 μM PSC833 (#SML0572, Sigma), respectively)^52,53^ and was supplemented with 5 μg/mL Hoechst 33342 to measure total nuclei. Cultures were incubated 15 min under an angle, in the dark at RT and subsequently acquired.

A z-stack imaging of the bottom layers of the culture is performed using the High-Content Confocal Imaging System. SUM projections were processed using Fiji Software. MF-SG and Hoechst 33342 intracellular fluorescence were quantified and subtracted for background signal. The relative intensity was calculated by normalizing substrate intensity to the nuclei count.

### Statistical data analysis

All statistical analysis were performed using Prism 9 Software (GraphPad). Data are presented as mean ± standard deviation (SD) or mean ± standard error of mean (SEM) when specified. Two-tailed unpaired t-tests were performed to determine the significance between two independent groups. One-way analyses of variance (ANOVA), followed by Tukey’s multiple comparisons test were performed when more than two groups were analyzed. Two-way ANOVA, followed by Tukey’s multiple comparison, was performed when examining the influence of 2 independent variables. For multiple comparison tests, all groups were compared between each other’s. Differences with p < 0.05 were considered as significant. *P < 0.05, **P < 0.01, ***P < 0.001 and ****P < 0.0001.

## Results

### Establishment of a multicellular placenta barrier on-a-chip with a physiologically relevant maternal-fetal interface

The maternal-fetal interface was established in the 3-lane OrganoPlate by co-culturing the human trophoblast cell line BeWo 30 and HUVEC (human umbilical vein endothelial cells) against a collagen-based scaffold (Figure 1). Phase contrast images showed that the BeWo b30 cells formed an epithelial barrier against the collagen-I/IV ECM gel after 3 days of culture, which progressed over time towards the vessel lane (Figure 2A). Interestingly, BeWo b30 in co-culture with HUVEC (Figure 2Aii) supported the maintenance of distinct maternal and fetal compartments. This compartmentalization was visualized by staining the epithelial maternal compartment for adherent junctions (ECAD), and both the epithelium and endothelium for Actin (Figure 2B). A 3D reconstruction of the placenta barrier model stained for Actin and tight junction marker ZO-1 (Figure 2C) showed two lumenized and perfusable tubules, providing access to their apical and basal sides. The placenta barrier co-cultures were highly reproducible across the platform (Figure 2D). During gestation, placenta barrier thickness is known to decrease, lowering to distances ranging between 20 μm and 100 μm in the first trimester of pregnancy^54,55^. The maternal-fetal interface distance of the placenta barrier on-a-chip culture was assessed after 9 days of culture by measuring the shortest, longest, and averaged distance between the epithelium and endothelium (Figure 2E). The maternal-fetal interface distance ranged from 13 to 119 μm in 75% of the cultures, with 25% below 49μm. The longest distance and average distance obtained ranges between 31-159 μm and 21-143 μm, respectively for 75% of the cultures. Those variable distances are comparable to the first-trimester *in vivo* situation.

**Figure 2.**
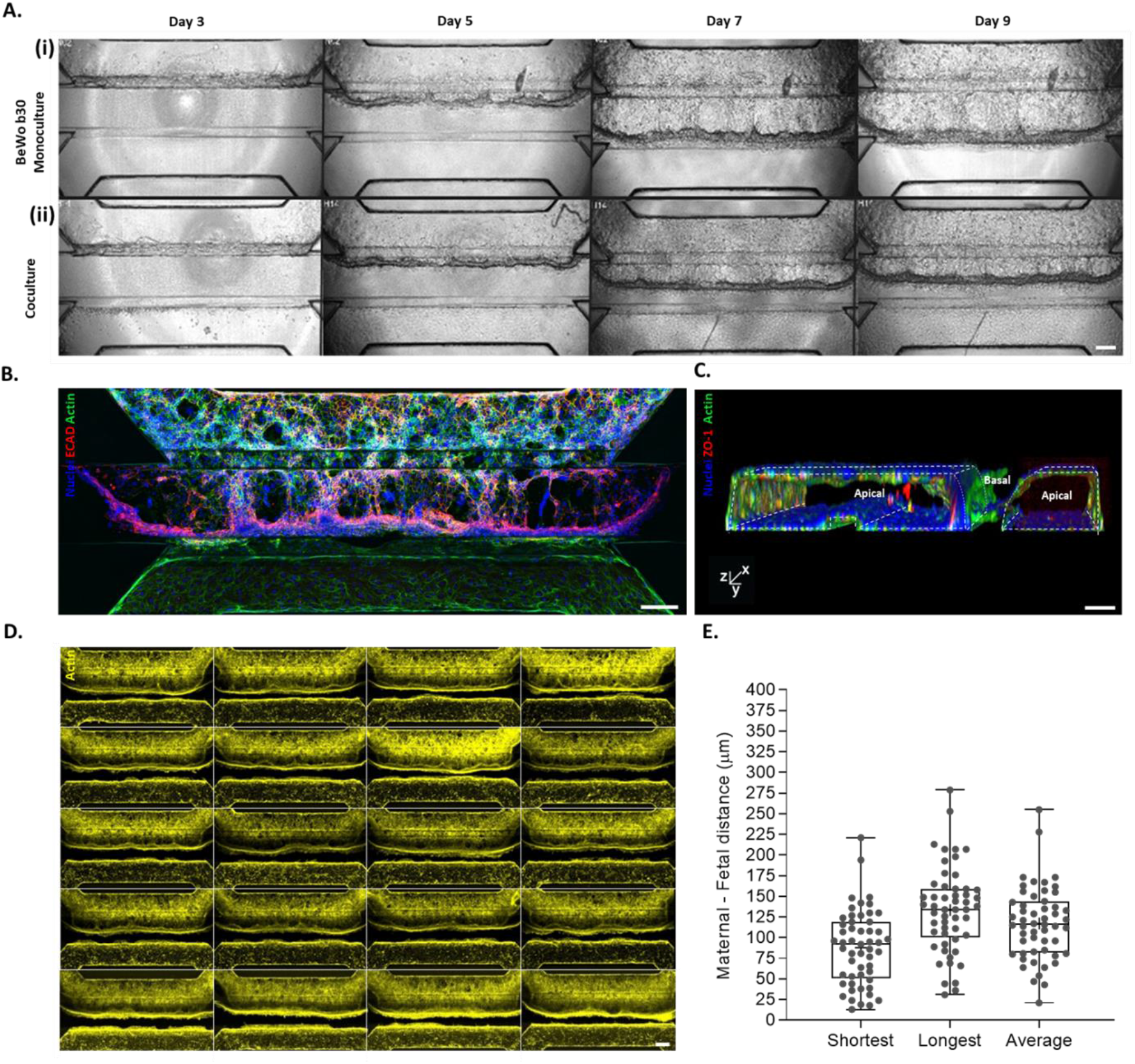
ECM thickness mimics the physiological placenta barrier interface in the early stage of pregnancy. (**A**) Representative phase contrast images of **(i)** BeWo b30 monoculture (maternal compartment), or **(ii)** co-culture with HUVEC (fetal compartment) depicting cell culture growth from day 3 to day 9. Middle channel contains a collagen-I/IV ECM gel. Scale bar: 200 μm. (**B**) Maximum intensity z-projection of the placenta barrier co-culture after 9 days of culture. Staining for Actin (green) and DNA (blue) shows the maternal-fetal interface, with the distinct maternal compartment stained for ECAD (red). Scale bar: 200 μm. (**C**) 3-dimensionnal reconstruction of the placenta barrier stained for ZO-1 (red), Actin (green) and DNA (blue) showing a lumenized maternal and fetal compartment in close proximity. (**D**) View of 20 chips containing placenta barrier cultures after 9 days of culture, stained for Actin (yellow). Scale bar: 100 μm. (**E**) Maternal-fetal interface distance quantification obtained from placenta barrier Actin immunostaining. For every chip, the shortest, longest, and average interface distance between mother and fetal compartment was measured. Data are plotted as box and whiskers (N=3, n=16-20). The line and plus sign represent the median and mean, respectively.

### Trophoblasts differentiate into a polarized syncytium with endocrine functions

*In vivo*, the syncytium is a polarized and secretory epithelium that is essential for the establishment and maintenance of pregnancy^56^. Those features were confirmed in the placenta barrier on-a-chip through gene and protein expression analysis. As shown by a positive immunostaining for microvilli marker Ezrin (Figure 3Ai) of the maternal compartment at 9 days of culture, the differentiated trophoblasts expressed microvillar proteins. The 3D reconstruction (Figure 3Aii), visualized by Actin and Ezrin, showed the formation of a confluent trophoblast layer with microvilli exclusively formed on the apical/luminal side of the epithelium. Syncytium formation was confirmed at the gene expression level (Figure 3B). After 48h of differentiation, BeWo b30 showed an up-regulation of syncytin-2 mRNA, a fusogenic marker involved in trophoblast fusion^57^. It was accompanied by an increase in hCG-α and hCG-ß gene expression, two sub-units of the human chorionic gonadotropin hormone, as well as PlGF, a placental growth factor. At the protein level, a staining of the epithelium for ZO-1 showed a breakdown of zona occludence-1 tight junctions (ZO-1) upon differentiation (Figure 3C). As visualized by nuclear and Actin staining, trophoblast differentiation led to the formation of a confluent layer of a mixed population of trophoblasts and multinucleated syncytiotrophoblasts. Moreover, a staining for hCG-ß and ECAD (Figure 3Di), as well as hCG-ß intensity quantification (Figure 3E) showed a significant increase of the hormone in the model, which was specifically produced within the syncytiotrophoblasts (Figure 3Dii). Additionally, while the hormone secretion in the lumen of the maternal compartment (Figure 3F) remained low and stable overtime (ranging between 497.2±102.1 pg/mL and 1005.1±307.9 pg/mL) in the undifferentiated culture, a significant time-dependent increase was observed upon differentiation, with a 4.5-fold (2145.7±307.9pg/mL) and 12.1-fold increase (5825.9±664.5pg/mL) (Figure 3G), after 24h and 48h differentiation respectively, compared to the undifferentiated epithelium.

**Figure 3.**
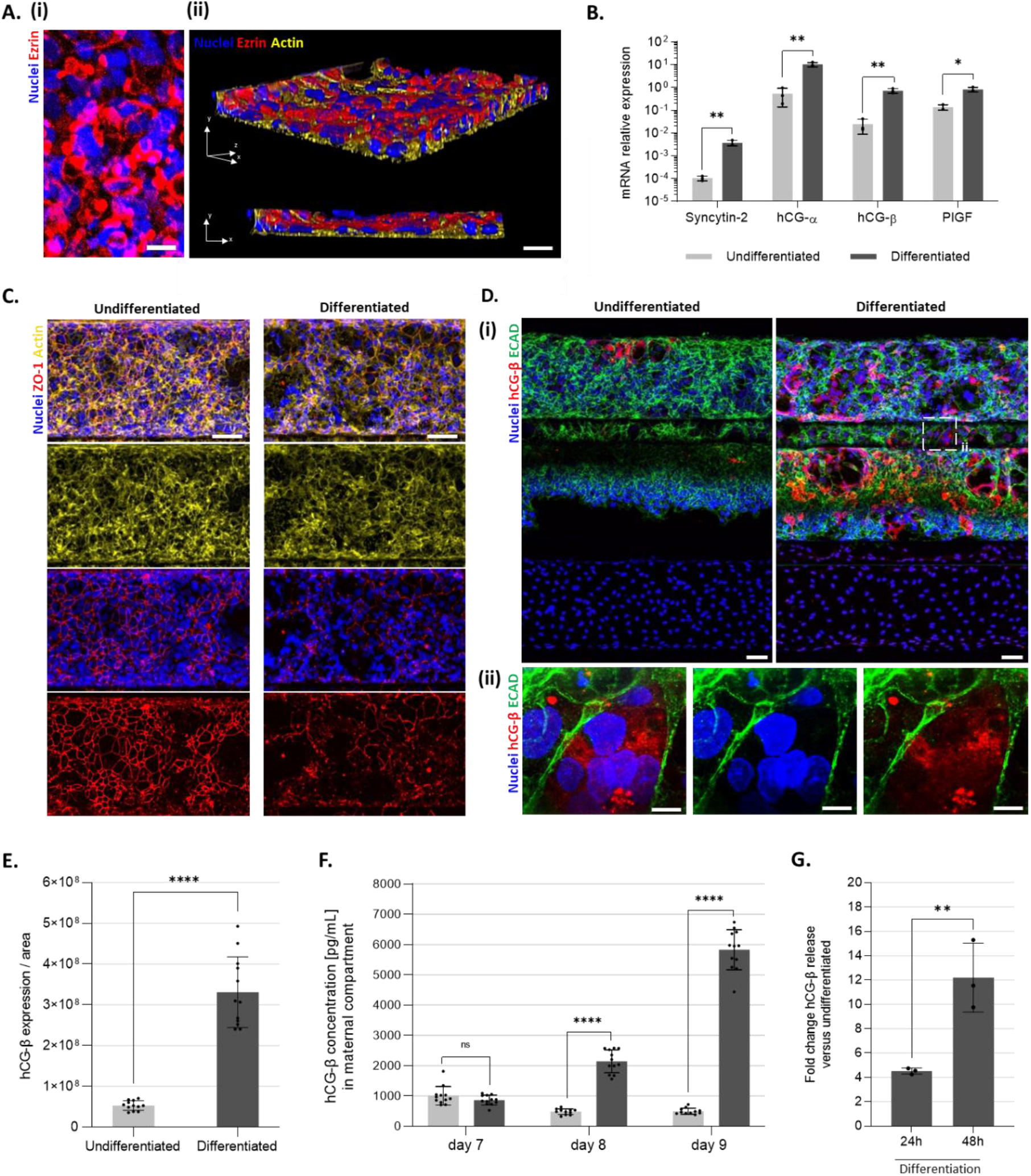
Trophoblasts differentiate into a polarized syncytium. (**A**) **(i)** Representative maximum intensity z-projection of syncytium microvilli stained for Ezrin (red) and DNA (blue). Scale bar: 25 μm. **(ii)** 3-dimensional reconstruction of the syncytium layer stained for Ezrin (red), Actin (yellow) and DNA (blue) shows a polarized syncytium. Scale bar: 50μm (**B**) mRNA expression of trophoblast differentiation markers Syncytin-2, hCG-α, hCG-ß and PlGF in undifferentiated or differentiated BeWo b30 (48h 25 μM Forskolin + 10 nM PMA). Each data point is derived from 5 pooled chips. Graphs show the mean of relative expression values normalized to GAPDH ± SEM and statistical analysis was performed using a one-way ANOVA (N=1, n=3). (**C**) Representative maximum intensity z-projection of undifferentiated and differentiated BeWo b30 stained for ZO-1 (red), Actin (yellow) and DNA (blue) shows the formation of multinucleated syncytium upon differentiation. Scale bar: 100μm (**D**) Representative and **(i)** maximum intensity z-projection of undifferentiated and differentiated BeWo b30 stained for ECAD (green), hCG-ß (red) and DNA (blue). Scale bar: 100μm. A zoom-in **(ii)** shows trophoblast hormone production within the formed syncytium. Scale bar: 20 μm. (**E)** Fluorescence intensity analysis of SUM projections of hCG-ß immunostaining from undifferentiated or differentiated BeWo b30. The intensity is corrected for the isotype control. Data points represent individual chips. Data was presented as mean ± standard deviation (SD) and statistical analysis was performed using two-tailed unpaired t-test (N=3, n=4). **(F)** Apical secretion of hCG-ß from undifferentiated and differentiated epithelium at day 7 (before differentiation), day 8 (24h differentiation) and day 9 (48h differentiation). Data points represent individual chips. Data was presented as mean ± SD and statistical analysis was performed using ordinary two-way ANOVA (N=3, n=4). **(G)** Fold change of hCG-ß release in the lumen of differentiated over undifferentiated BeWo b30 at day 8 (24h differentiation) and day 9 (48h differentiation). Data points represent the average of hCG-β concentration of differentiated over undifferentiated epithelium of 4 chips. Data was acquired from 3 individual experiments. Data were presented as mean ± SD and statistical analysis was performed using two-tailed unpaired t-test (N=3, n=4). ns: non-significant; *P < 0.05; **P < 0.01; ****P < 0.0001.

### Placenta barrier on-a-chip is suitable for paracellular and drug transport studies

In order to provide a tool for transport assessment across the maternal-fetal interface, the placenta barrier on-a-chip model was characterized for basement membrane production, crucial for solute filtration, as well as for barrier integrity and drug transport activity. To verify the molecular composition of the basement membrane, syncytium and endothelium were stained for diverse basement membrane proteins (Figure 4A). Both maternal and fetal compartments secreted Laminin and Perlecan (Figure 4Ai), Collagen-IV (Figure 4Aii) and Nidogen-2 (Figure 4Aiii). In contrast, only the endothelium produced Fibronectin (Figure 4Aii), while Tenascin-C was solely secreted by the syncytium (Figure 4Aiii). The syncytium constitutes the main physical barrier for paracellular transport of solutes^55^. To characterize syncytium barrier tightness in our model, a fluorescent 4.4kDa TRITC-labelled dextran was administrated to the lumen of the trophoblast tubule (Figure 4B). Leakage from the lumen to the adjacent gel compartment was monitored for undifferentiated and differentiated trophoblast tubules at 7, 8 and 9 days of culture. Both undifferentiated and differentiated epithelium formed a leak-tight barrier that was stable over time, showing low and non-significant leakage scores ranging from 0.19±0.10 and 0.17±0.11 on day 7 to 0.08±0.03 and 0.10±0.02 on day 9, for undifferentiated and differentiated epithelium, respectively (Figure 4C). In the same manner, fluorescein (0.376 kDa) transfer across the trophoblast barrier was assessed from days 7 to 9 (Supplementary figure 1). At 9 days of culture, fluorescein paracellular transport (0.10±0.02 leakage score) is increased by 38.5% compared to 4.4kDa TRITC-labelled Dextran (0.13±0.03 leakage score) (Figure 4D). The transepithelial electrical resistance (TEER) (Figure 4E) revealed that an undifferentiated epithelium barrier increased its tightness over time with TEER values ranging from 7.56±1.83 kΩ on day 6 to 13.50±1.35 kΩ on day 9, whereas trophoblast epithelium showed a time-dependent decrease upon differentiation with TEER values dropping from 13.19±2.24 kΩ on day 7 to 8.17±1.71 kΩ on day 9.

**Figure 4.**
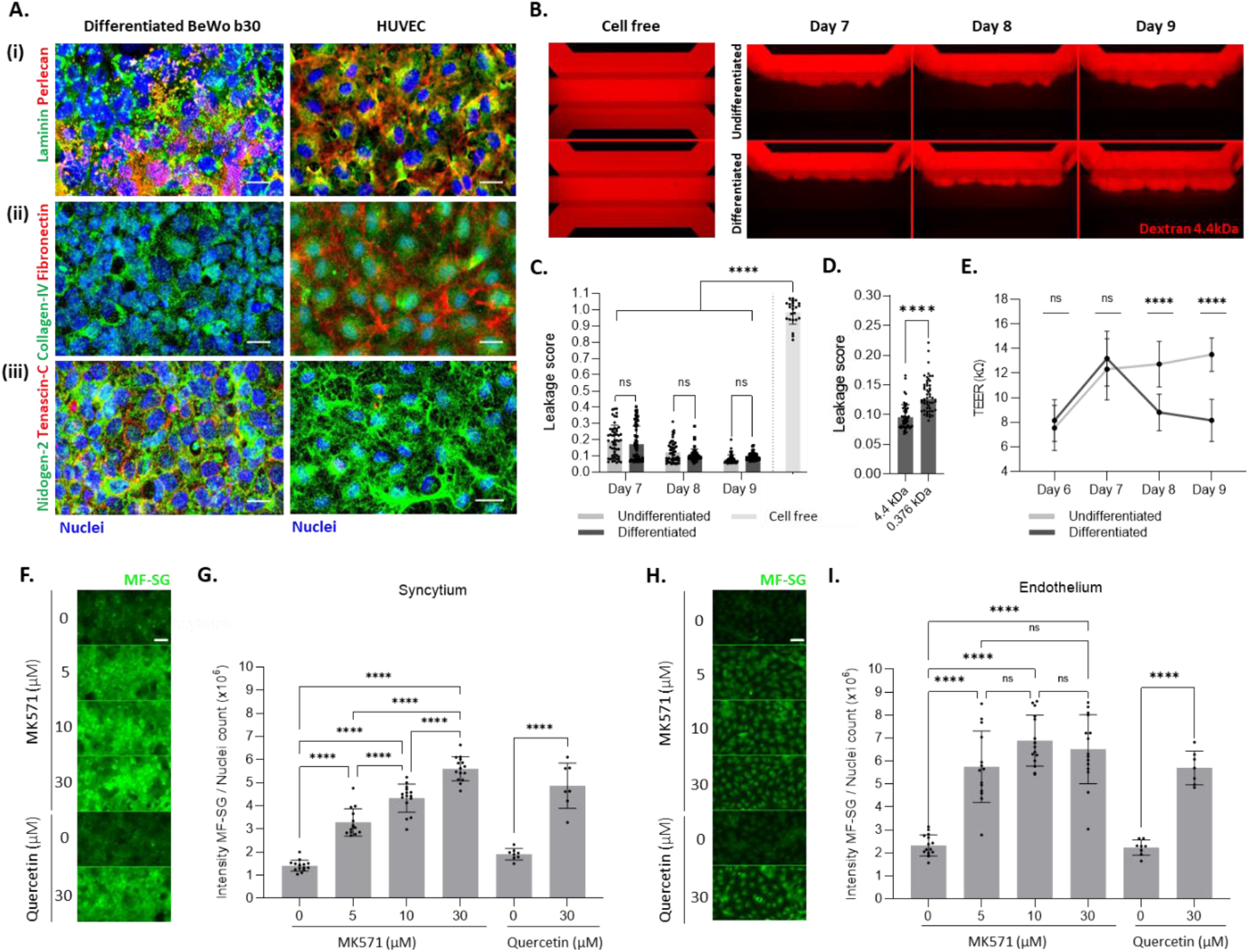
Placenta on-a-chip forms a leaktight barrier and is suitable for drug transport studies. **(A)** Representative maximum intensity z-projection of syncytium and endothelium stained for DNA (blue) and the basement membrane proteins **(i)** laminin (green) and perlecan (red), **(ii)** collagen-IV (green) and fibronectin (red), **(iii)** nidogen-2 (green) and tenascin-C (red). **(B)** Fluorescent images of representative undifferentiated and differentiated placenta barrier co-culture, perfused in the maternal compartment with fluorescent molecules (red) using 4.4 kDa TRITC-Dextran for 12 min on days 7, 8 and 9. Trophoblast differentiation was started on day 7 in the Differentiated condition. The cell free control showed passage of the fluorescent dye into the adjacent gel and perfusion channel, whereas fluorescent dye was retained by undifferentiated and differentiated trophoblast tubules. **(C)** Ratio of 4.4kDa TRITC-Dextran intensity in the basal side of the maternal tube versus the intensity at the apical side of the maternal tube (Leakage score) after T=12min, on days 7, 8 and 9. Data points represent individual chips. Data are presented as mean ± SD and statistical analysis was performed using ordinary two-way ANOVA (N=3, n=17-19 for undifferentiated and differentiated conditions; N=3, n=6-9 for cell free) **(D)** Ratio of 4.4 kDa TRITC-Dextran and Fluorescein (0.376 kDa) intensity in the basal side of the maternal tube versus the intensity at the apical side of the maternal tube (Leakage score) in differentiated trophoblast at 9 days of culture after T= 12min. Data are presented as mean ± SD and statistical analysis was performed using two-tailed unpaired t-test (N=3, n=17-18). **(E)** Transepithelial electrical resistance (TEER) measured on undifferentiated and differentiated trophoblasts on days 6, 7, 8 and 9. Trophoblast differentiation was started on day 7 in the Differentiated condition. Data is presented as mean ± SD and statistical analysis was performed using ordinary two-way ANOVA (N=3, n=9-21). **(F)-(I)** Evaluation of Multi-Drug Resistance Protein (MRP) transporter activity in the placenta barrier on-a-chip. The non-fluorescent CMFDA was used as MRP substrate. It enters the cells passively and transforms inside the cells into its fluorescent product MF-SG. **(F)** & **(H)** Representative summary z-projections of intracellular GS-MF (green) intensity in differentiated trophoblasts (F) and endothelium (H) treated for 30 min with vehicle (DMSO) or two MRP specific inhibitors, MK571 (5 μM, 10 μM, 30 μM) and quercetin (30 μM). **(G) & (I)** Ratio of GS-MF intensity over nuclei count in the syncytium (G) and endothelium (I). A dose-dependent inhibition of MRP transporters by MK571 and Quercetin is observed. Data points represent individual chips. Data are presented as mean ± SD and statistical analysis was performed using ordinary one-way ANOVA (N=3, n=4 for MK571; N=2, n=4 for Quercetin). ns: non-significant; ****P < 0.0001.

Throughout pregnancy, drug transporters in the placenta protect the fetus from xenobiotics^58^. We therefore demonstrated the functionality of the multi-drug resistance protein (MRP) family in our placenta model. The model expressed MRP1, MRP2, MRP5 and less abundantly MRP3 and MRP4 at the mRNA level (Supplementary figure 2). The cell tracker 5-chloromethylfluorescein diacetate (CMFDA) and its fluorescent metabolite MF-SG are well-known MRP substrates^59^. MRP-mediated transport was assessed on day 9 of culture in the syncytium and endothelium using MK571 and quercetin, two MRP inhibitors. In presence of MK571 and quercetin in both maternal (Figure 4F, G) and fetal (Figure 4H, I) compartments, there was a dose-dependent increase in the intracellular fluorescence intensity indicating the inhibition of MRP-mediated efflux transport in both syncytium and endothelium. These data confirmed MRP transporter activity in the placenta barrier model. In the same way, the Breast Cancer Resistance Protein (BCRP), which restricts the distribution of its substrates to the fetus, was active in the syncytium of the placenta barrier-on-a-chip setup (Supplementary figure 3).

### Impaired oxygen tension and perfusion induces molecular and functional changes in the placenta barrier that closely resembles those observed in preeclampsia

Hypoxia and ischemia are central mechanisms of damage to the placenta leading to induction of preeclampsia (PE)^60^. In order to model preeclamptic features in the placenta barrier on-a-chip model, cultures were exposed to a combination of different oxygen and perfusion tensions. Parameters to mimic healthy conditions included 20% O_2_ and perfusion flow and are here referred to as the “Normoxic” condition (Figure 5Ai). Preeclamptic characteristics were mimicked by either switching the culture to 1% O_2_ with medium perfusion flow for 24h on day 8 (“Hypoxic”) or to 1% O_2_ and static medium conditions (“Ischemic”). Phase contrast images of the placenta barrier did not show a change in syncytium integrity after 24h reduction of low O_2_ tension (“Hypoxic”; Figure 5Bi) compared to the “Normoxic” condition (Figure 5Bii). However, syncytium barrier damage could be observed in the low oxygen and ischemic environment, as shown by darker cells (“Ischemic”; Figure 5Biii). Syncytium damage was confirmed by measuring the transepithelial electrical resistance (TEER) through the syncytium (Figure 5C) on day 8 and day 9. Both “Normoxic” and “Hypoxic” conditions showed respectively a 1.23 and 1.29-fold decrease in TEER values over time, indicating hypoxia itself did not affect the epithelium barrier. In contrast, “Ischemic” condition triggered a significant 1.9-fold change decrease in syncytium permeability compared to the “Normoxic” control on day 9, correlating with cell damage observed in the phase contrast images. Interestingly, nuclei count over a representative syncytium area (Supplementary figure 4A) showed 10% more cells in the “Ischemic” condition (Figure 5D, 5E) compared to the “Normoxic” condition, an increase which was not observed in the “Hypoxic” condition.

**Figure 5.**
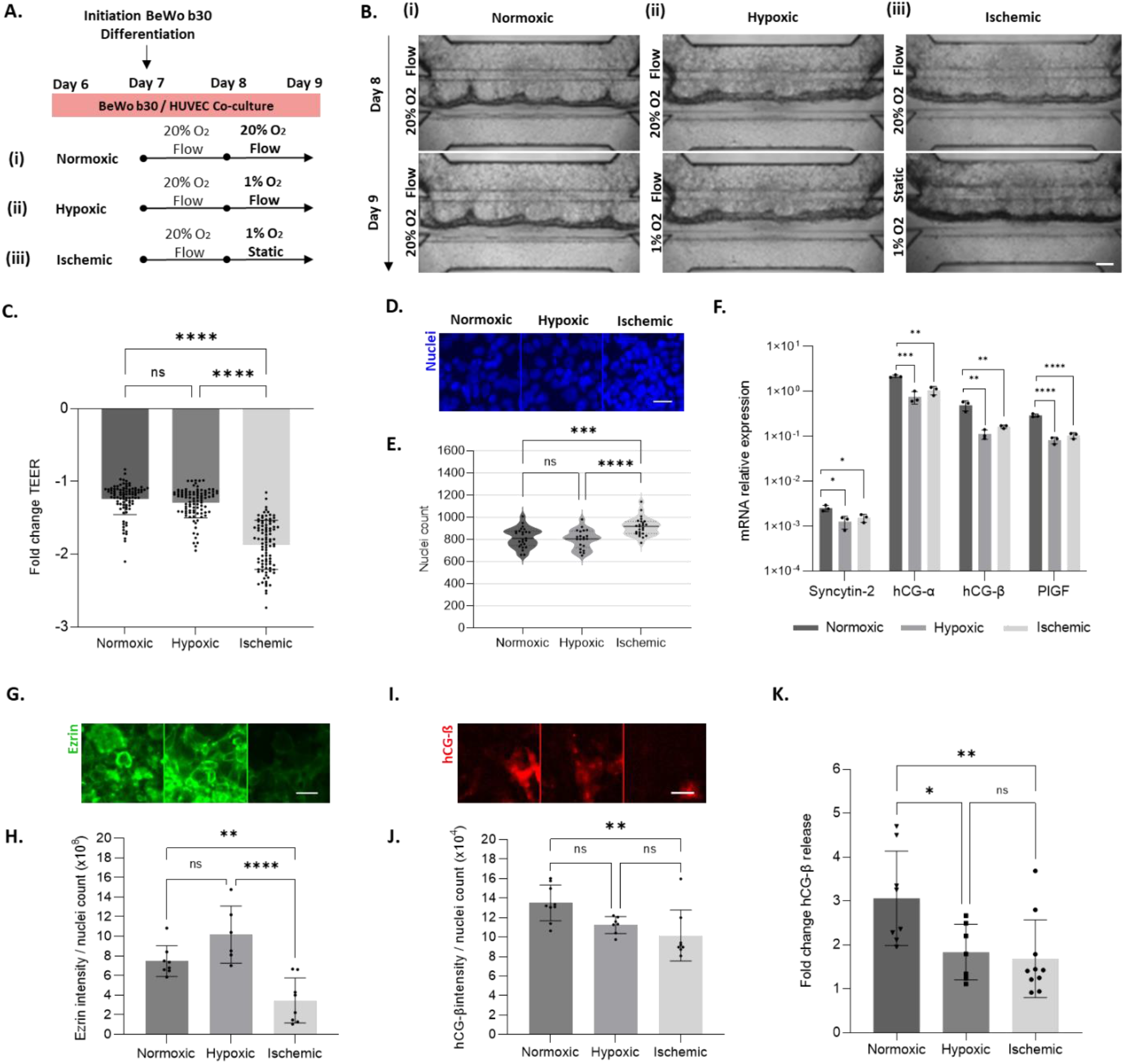
Low oxygen tension and lack of perfusion result in syncytium damages resembling preeclamptic characteristics. (**A)** Workflow depicting the parameters to mimic healthy and preeclamptic conditions. On day 8, the placenta barrier model is either **(i)** kept under standard condition: 20% O2 and flow (“Normoxic”), **(ii)** cultured for 24h under 1% O2 and flow (“Hypoxic”) or **(iii)** cultured for 24h under 1% O2 and static (“Ischemic”). (**B)** Representative phase contrast images of the placenta barrier on-a-chip model depicting **(i)** “Normoxic”, **(ii)** “Hypoxic” and **(iii)** “Ischemic” conditions over 8 and 9 days of culture. Scale bar: 200μm. (**C)** Fold change decrease of transepithelial electrical resistance (TEER) values of the syncytium on day 9 versus day 8 in “Normoxic”, “Hypoxic” and “Ischemic” conditions. Data points represent individual chips. Data are presented as mean ± SD and statistical analysis was performed using ordinary one-way ANOVA (N=3, n=30-33). (**D)** Representative summary z-projections of syncytium nuclei count in “Normoxic”, “Hypoxic” and “Ischemic” conditions after 9 days of culture. **(E)** Quantification of the nuclei count in “Normoxic”, “Hypoxic” and “Ischemic”. Data points represent individual chips. Data points are represented as dispersion, with the average (solid line) ± SD (dotted line) and statistical analysis was performed using ordinary one-way ANOVA. (N=3, n=21-25). **(F)** mRNA expression of trophoblast differentiation markers Syncytin-2, hCG-α, hCG-ß and PlGF in “Normoxic”, “Hypoxic” and “Ischemic”. Each data point is derived from 5 pooled chips. Graphs show mean of relative expression values normalized to Actin ± SEM and statistical analysis was performed using ordinary one-way ANOVA (N=1, n=3). **(G)** Representative summary z-projections of Ezrin intensity in “Normoxic”, “Hypoxic” and “Ischemic” conditions. Scale bar: 50μm. **(H)** Ratio of Ezrin intensity over nuclei count of the syncytium, in “Normoxic”, “Hypoxic” and “Ischemic” conditions. Data points represent individual chips. Data are presented as average signal intensity / nuclei count ± SD and statistical analysis was performed using ordinary one-way ANOVA (N=2-3, n=2-3). **(I)** Representative summary z-projections of hCG-ß intensity in “Normoxic”, “Hypoxic” and “Ischemic” conditions. Scale bar: 50μm. **(J)** Ratio of hCG-ß intensity over nuclei count in the syncytium, in “Normoxic”, “Hypoxic” and “Ischemic” conditions. Data points represent individual chips. Data are presented as mean ± SD and statistical analysis was performed using ordinary one-way ANOVA (N=3, n=2-3). **(K)** Fold change increase of apical secretion of hCG-ß into the maternal compartment lumen in “Normoxic”, “Hypoxic” and “Ischemic” conditions on day 9 versus day 8. Data points represent individual chips. Data are represented as mean ± SD and statistical analysis was performed using ordinary two-way ANOVA (N=3, n=3-4). ns: non-significant, *P < 0.05; **P < 0.01; ***P < 0.001; ****P < 0.0001).

The different syncytia were then characterized at the gene expression level (Figure 5F) for markers known to be mis-regulated during PE. In comparison to the “Normoxic” model, the fusogenic marker, syncytin-2 exhibited a 1.58-fold decrease in the “Hypoxic” and 0.58-fold change decrease in the “Ischemic” conditions. Likewise, the mRNA level of two isoforms hCG-α and hCG-β were downregulated to a greater extent in “Hypoxic” with a 1.52-fold change and 2.13-fold change respectively, compared to 1.04-fold change and 1.57-fold change for “Ischemic” conditions. Finally, placental growth factor PlGF, a predictive biomarker of preeclampsia^61^ also showed a decrease in expression under low oxygen tension with a 1.81-fold change decrease, while it was reduced to a 1.48-fold change after exposure to hypoxia-ischemia environment.

The syncytium was further characterized at the protein level. Immunostaining for the microvilli marker, Ezrin (Figure 5G) showed a modest increase in the “Hypoxic” condition compared to the “Normoxic” placenta model that was not statistically significant (Figure 5H). However, the “Ischemic” condition exhibited 50% lower expression of Ezrin than the “Normoxic” placenta model. Moreover, patched loss of Ezrin signal in the “Ischemic” condition showed a complete loss of microvilli in those cell surface areas (Supplementary figure S4B). Compared to the “Normoxic” condition, the signal for hCG-β (Figure 5I, 5J) was decreased for the “Hypoxic” condition, which was amplified to 25% for the “Ischemic” condition. This reduction in hCG-β was confirmed at the secretion level by measuring the hormone release into the syncytium lumen (Figure 5K) before and after 24h exposure to the different oxygen/perfusion environments. Indeed, while the hormone showed a 3.06-fold increase in secretion in the “Normoxic” placenta model, only a 1.84-fold and 1.67-fold increase was observed in “Hypoxic” and “Ischemic” respectively. Additionally, glucose transporter-1 (GLUT-1), a major regulatory factor in the process of maternal-fetal glucose exchange was evaluated at the protein level expression (Supplementary figure 5A,B) and demonstrated to decrease 25% after 24h culture under “Ischemic” conditions compared to the control. Taken together, these data showed that hypoxia itself or combined with static conditions could induce phenotypical changes in our placenta barrier co-culture.

## Discussion & Conclusions

In this study, we report *in vitro* modeling of preeclampsia (PE) characteristics in the form of a placenta barrier on-a-chip with a pathology-associated phenotype in a high-throughput microfluidic setup. The healthy placenta barrier model recapitulated essential features of early pregnancy, such as barrier formation, polarization, transporter function and hormone secretion. Exposure of the placenta barrier model to an ischemic and hypoxic environment triggered alterations in syncytium integrity and functionality at different degrees, representative of syncytial alterations linked to PE.

Progressing with days in culture, trophoblasts exhibited invasive behavior (Figure 2) likely supported by the secretion of a wide range of metalloproteinases (MMPs)^42^ and characteristics of their malignant properties. While an MMP-8 inhibitor was used during the differentiation process to target collagen-I digestion^43^, the fetal endothelium proved important for the stabilization of the materno-fetal distance. The placenta barrier interface distance ranged between 21-143 μm, with variations within the same chip, which is in line with studies reporting an irregular maternal-fetal diffusion distance of 20-100 μm in the first trimester of pregnancy *in vivo*^55,62^. The co-culture set-up of ECM and cells in a membrane-free manner provides a unique advantage for modeling this progressive thinning of the maternal interface that would not be present for membrane-based systems such as Transwell or microfluidics devices that make use of artificial membranes ^25,32^.

Characterization of the differentiated syncytia demonstrated the establishment of apicobasal polarization (Figure 3), with microvilli densely covering the apical side of the maternal epithelium, reflecting the *in vivo* syncytium with an extensive microvillous brush barrier in contact with the maternal blood^63^. These microvilli are essential to increase the available surface area for material exchange and transporter polarization *in vivo*^64,65^. The 48h differentiation of BeWo b30 resulted in a significant up-regulation of syncytin-2, a gene playing a critical role in the fusion of trophoblasts^41^. It was translated at the molecular level by a loss of ZO-1 tight junctions, leading to the formation of an epithelium consisting of a mixed population of trophoblasts and multinucleated syncytiotrophoblasts, characteristics of the first-trimester pregnancy syncytium^35^. Up-regulation of hCG subunits was confirmed at the mRNA level and hCG-ß production was significantly increased and secreted into the maternal compartment. These data are in agreement with previous studies in BeWo cell lines reporting elevated hormone release upon chemical stimulation with Forskolin^66,67^. *In vivo*, hCG-ß is regarded as one of the main markers of differentiation and its increased production in the first months of gestation is critical for the maintenance of early pregnancy^41^. Finally, syncytium formation was associated with an elevation of placental growth factor (PlGF) mRNA, a proangiogenic factor known to mainly be produced by syncytiotrophoblasts with the role to promote placental angiogenesis during pregnancy^68^. Taken together, these data confirmed a successful syncytium development in the placenta barrier co-culture.

On their basal sides, the syncytium and endothelium successfully secreted their own basal lamina as observed by the production of laminin, heparan sulfate (perlecan), collagen-IV and nidogen-2 (Figure 4). The trophoblasts also expressed tenascin-C, a matrix protein involved in syncytial fusion and playing a critical role in placental homeostasis development^69^. Syncytium did not secrete fibronectin, which may contribute to their malignant origin^70^. Those secreted proteins are well known to act both as functional support and as a filtration barrier between the maternal-fetal interface^71^. Assessment of the paracellular flux of two different molecular-weight solutes across the maternal barrier demonstrated the formation of a leaktight syncytium for 4.4kDa molecules (Figure 4C), with a higher permeability to fluorescein (0.376 kDa) (Figure 4D) increasing upon differentiation (Supplementary figure 1). Those results corroborate those previously reported, demonstrating a size-dependent passive diffusion of molecules across BeWo cells and primary trophoblasts *in vitro*^72^ and a passive diffusion of drugs <500 Da across the syncytium *in vivo*^54^. Similarly, the absolute TEER (Ω) values, calculated to discriminate the impact of epithelium changing surface upon differentiation, showed a decrease upon syncytium formation (Figure 4E). Additionally, the placenta barrier on-a-chip presents active MRP and BCRP drug transporters. The combination of CMFDA substrate with MK175 or quercetin suggested that MRP1, the most prominently expressed transporter subtype in cytotrophoblasts and syncytiotrophoblasts of first trimester placenta^73^, was active^54,74^. We thus effectively developed *in vivo-*like features making our placenta barrier on-a-chip a relevant tool for transplacental transport studies.

Insufficient uteroplacental oxygenation together with ischemia is considered to be among the leading causes of PE^75^. We mimicked the preeclamptic environment by exposing the placenta barrier-on-a-chip to 1% O_2_ level, either under perfusion (“Hypoxic”) or static (“Ischemic”) conditions, for 24 hours (Figure 5). While hypoxia itself did not cause syncytium disruption, removal of perfusion flow resulted in a significant increase of permeability which was associated with a loss of syncytial microvilli. Those results are in line with studies showing that induced-hypoxia in static BeWo b30 cultures was accompanied by defective paracellular transport through alteration of tight-junctions^76^. While there is limited *in vivo* data available regarding syncytium permeability during PE, the pathology is known to be associated with syncytial degeneration, damaged syncytium integrity and loss of microvilli^35,77^, strongly suggesting alteration in syncytium permeability^78^. Syncytium damages observed under ischemia were likely triggered by a loss of flow, reducing diffusion of nutrients and removal of waste in the placenta barrier on-a-chip. The *in vitro* “Ischemic” model showed an increase in the trophoblast cell population, in line with an *in vivo* comparative study in placenta issued from normal and preeclamptic pregnancy revealing a higher proliferation rate of villous trophoblast in the diseased placenta^79,80^. While some designated these excessive proliferative cells as immature and intermediate trophoblasts^80^, others hypothesized that this higher proliferation rate is a mechanism to counteract syncytium damage^35^.

A down-regulation of markers involved in syncytium differentiation, such as Syncytin-2, hCG-α, hCG-ß, was observed at the gene expression level in both preeclamptic models. The loss of syncytin-2 is in line with earlier studies showing that PE-mimicking hypoxia in BeWo b30 and primary trophoblast cells^81,82^ or cells isolated from PE placentas^83^ presented reduced cell fusion. Moreover, another study stipulated that syncytin downregulation correlates with the severity of PE, and can serve as a marker to evaluate women at risk of developing PE during pregnancy^84^. Regarding hCG-ß, the downregulation at the gene expression level was translated at the molecular level by a reduced hormone production and secretion into the maternal compartment, with a more pronounced effect when low oxygen tension was combined with the static condition (Figure 5K). This correlates with the evidence that low hCG-ßconcentrations late in the first trimester may be associated with a subsequent risk of preeclampsia.^85^ Also, low PlGF expression and concentration observed in preeclampsia, a biomarker of PE pathology^61^, was replicated at the mRNA level in our “Hypoxic” and “Ischemic” models.

The trophoblast cell line used in our model has been widely used in placental research. It has been reported to display several morphological properties and functions common to primary trophoblasts, such as differentiation into syncytiotrophoblasts, hormones production and polarized transcellular transports^72^. However, due to their choriocarcinoma origin, they do not differentiate spontaneously in vitro^86^ and show different methylation patterns compared to primary trophoblasts, contributing to variation in the expression of numerous genes, imply that care must be taken when extrapolating results^87^. While differences are observed in placental drug transporter expression^88^, further studies assessing transplacental transfer of drugs by mass spectrometry would increase insight in transport capabilities of our model. In the past year, tremendous efforts have been conducted in the development of more physiologically relevant alternatives with the generation of cytotrophoblast stem-like cells from hPSCs^89,90^ or long-term trophoblast organoids^91^. Their integration into our platform would move the placenta barrier on-a-chip a step further in the modeling and study of normal and diseased placenta in early pregnancy^89^. Regarding the oxygen tension used in the co-culture set-up, the 20% O_2_ level differs from the hypoxic placental environment which reaches up to 6-8% at the end of the first trimester^7^. However, we choose to adhere to common practice to culture in a 20% O_2_ environment, as this is more practical for long-term culture, avoids oxygen adjustments and allows us to compare our results with data from the literature.

Lastly, PE is characterized by an intermittent maternal blood flow within the intervillous space associated with ischemia/hypoxia – reperfusion type of injury. While we only focused on the hypoxic and ischemic/hypoxic aspect of the disease, reperfusion modeling would be of a great interest, especially for the study of oxidative stress and release of anti-angiogenic factors, which are other hallmarks of PE.

In conclusion, we described here the first placenta barrier on-a-chip system modeling preeclamptic characteristics. The model recapitulates key features of the placenta barrier in early pregnancy such as syncytium differentiation, barrier function, polarization, hormone secretion and transporter abilities. In addition, we could effectively recapitulate hallmarks of preeclampsia such as reduced barrier function, hormonal secretion, brush border formation and nuclei count. The healthy model represents a powerful tool for modeling drug placental transfer assessment during pregnancy. Exposure to preeclamptic conditions will prove beneficial to improve our understanding of the pathology. Its potential for high-throughput screening approaches will open tremendous opportunities in the identification of novel targets and the development of novel therapies for the prevention or treatment of the disease.

## Author contributions

G. Rabussier, H.L. Lanz and K. Bircsak designed the study. G. Rabussier, I. Bünter and J. Bouwhuis performed experiments and data analysis. H.L. Lanz, K. Bircsak, K. Dormansky, C.E. Mudoch and L.J. de Windt supervised the research. G. Rabussier, H.L. Lanz and K.M. Bircsak wrote the paper. G. Rabussier, C. Soragni, C. Ping Ng, H.L. Lanz and K. Bircsak contributed to the visualization. All authors read and approved the final manuscript.

## Acknowledgements

We would like to acknowledge Frederik Schavemaker for the artwork.

## Supplementary Materials and methods

### BCRP activity assay

BCRP activity measurement was assessed as previously described for MRP transporters. 30uL of 7 μM Hoechst 33342 (BCRP substrate) diluted in Opti-HBSS (1:2 Opti-MEM and HBSS) was added to all the perfusion inlets and outlets, either with inhibitors (1μM, 10μM Ko143) or corresponding vehicle (DMSO). OrganoPlates were incubated in a humidified incubator (37°C, 5% CO_2_) for 30min on the OrganoFlow (7°, 8-min intervals). Cultures were then cooled down by adding 50uL cold HBSS in the observation window. Perfusion lanes assay solution was replaced with iced-cold inhibitor cocktail (10μM MK-571, 10μM Ko143, 10μM PSC833). Cultures were incubated 5min under an angle, in the dark at RT and subsequently acquired.

### Supplementary figures

**Supplementary figure 1.**
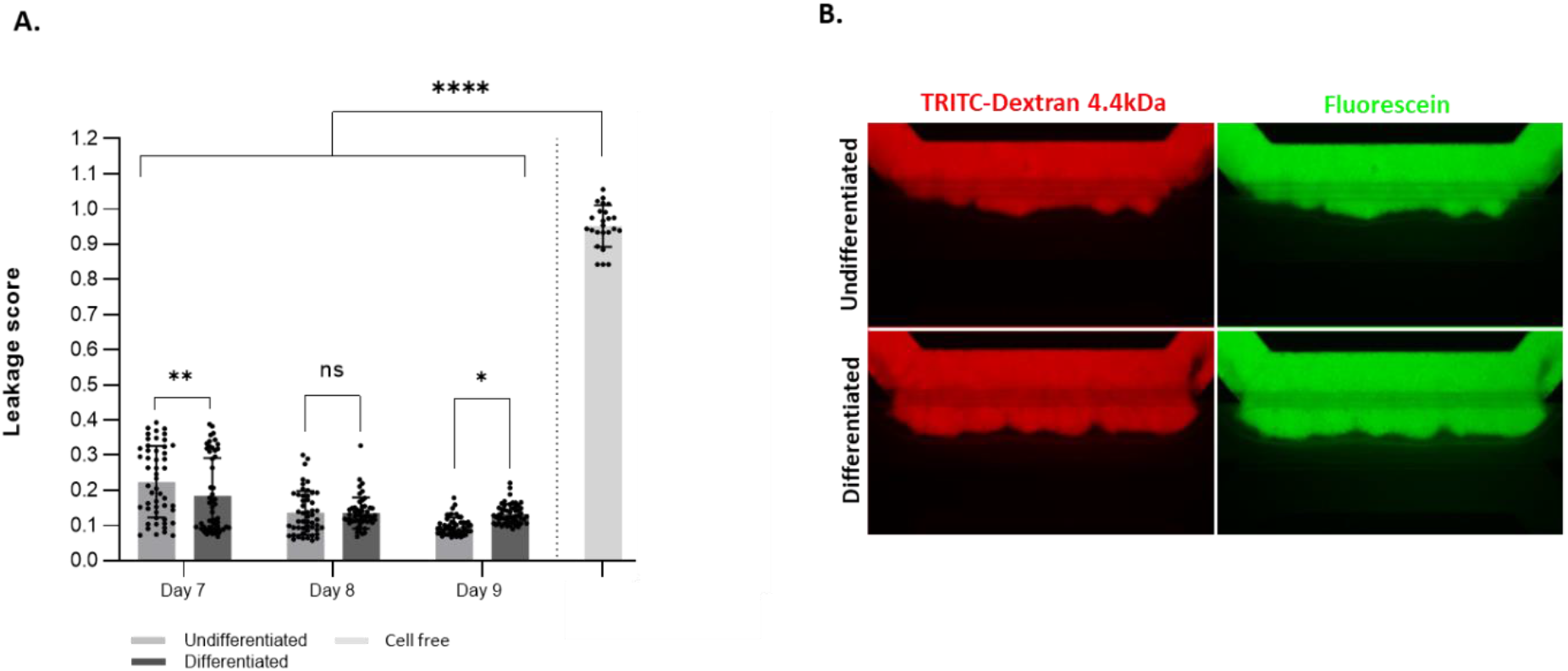
Placenta on-a-chip forms a leaktight barrier with increased permeability upon differentiation for low-molecular weight molecules. **(A)** Ratio of fluorescein (0.376 kDa) intensity in the basal side of the maternal tube over the intensity in the apical side of the maternal tube (Leakage score) after T=12min, on days 7, 8 and 9. Data points represent individual chips. Data are presented as mean ± SD. Statistical analysis was performed using two-way ANOVA. (ns: non-significant; *P < 0,05; **P < 0,01; ****P < 0,0001). (N=3, n=17-19 for undifferentiated and differentiated conditions; N=3, n=6-9 for cell free). **(B)** Fluorescent images of representative undifferentiated and differentiated placenta barrier co-culture at 9 days of culture, perfused in the top channel with TRITC-labelled Dextran (red) and fluorescein (green), after T=12 min.

**Supplementary figure 2.**
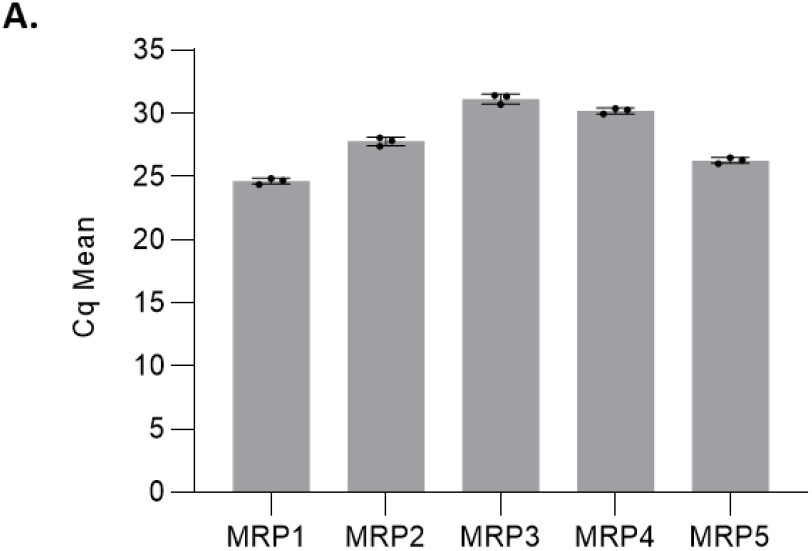
Differentiated BeWo b30 express Multidrug resistance protein (MRP) transporters. **(A)** Graph shows Cq mean values for MRP1, MRP2, MRP3, MRP4 and MRP5. Each data point is derived from 5 pooled chips. Data are presented as mean ± SEM. (N=1, n=3)

**Supplementary figure 3.**
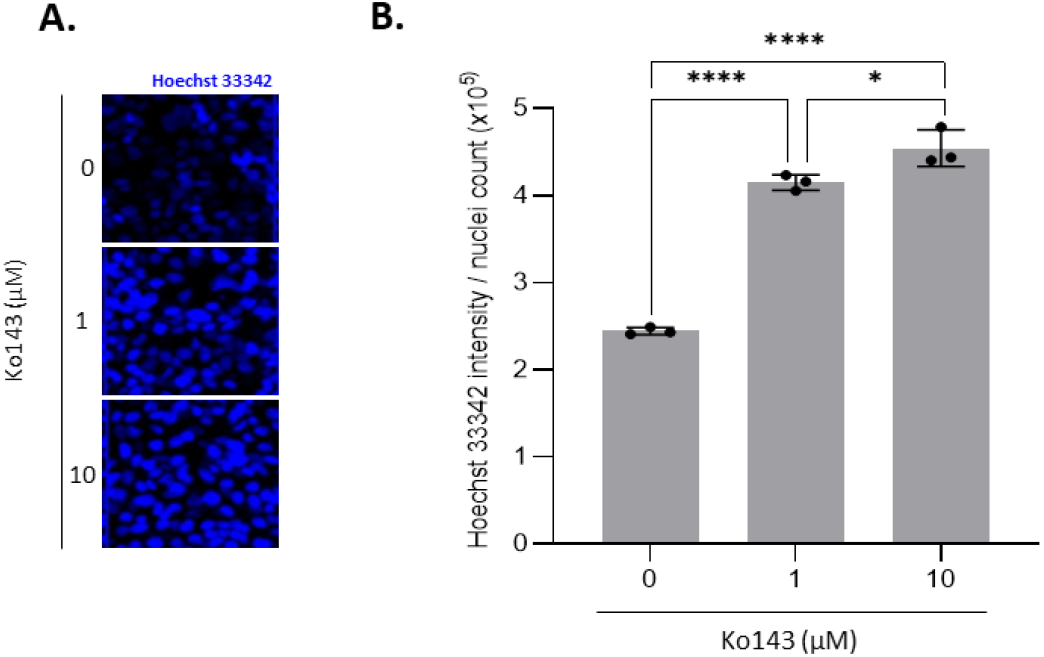
Evaluation of Breast Cancer Resistance Protein (BCRP) transporter efflux activity in the placenta barrier on-a-chip. Hoechst 33342, a nuclear-specific dye that binds to DNA is used as a substrate. It enters the cells passively and binds to DNA with an increased fluorescence **(A)** Representative summary z-projections of intracellular Hoechst 33342 (blue) intensity in differentiated trophoblasts untreated (Vehicle control) and treated for 30min with a BCRP inhibitor, Ko143 (1μM, 10μM). **(B)** Ratio of Hoechst 33342 intensity over nuclei count of the syncytium, in untreated or exposed culture to Ko143. A dose-dependent efflux inhibition by Ko143 is observed. Data points represent individual chips. Data are presented as mean ± SD and statistical analysis was performed using ordinary one-way ANOVA. (N=1, n=3).

**Supplementary figure 4.**
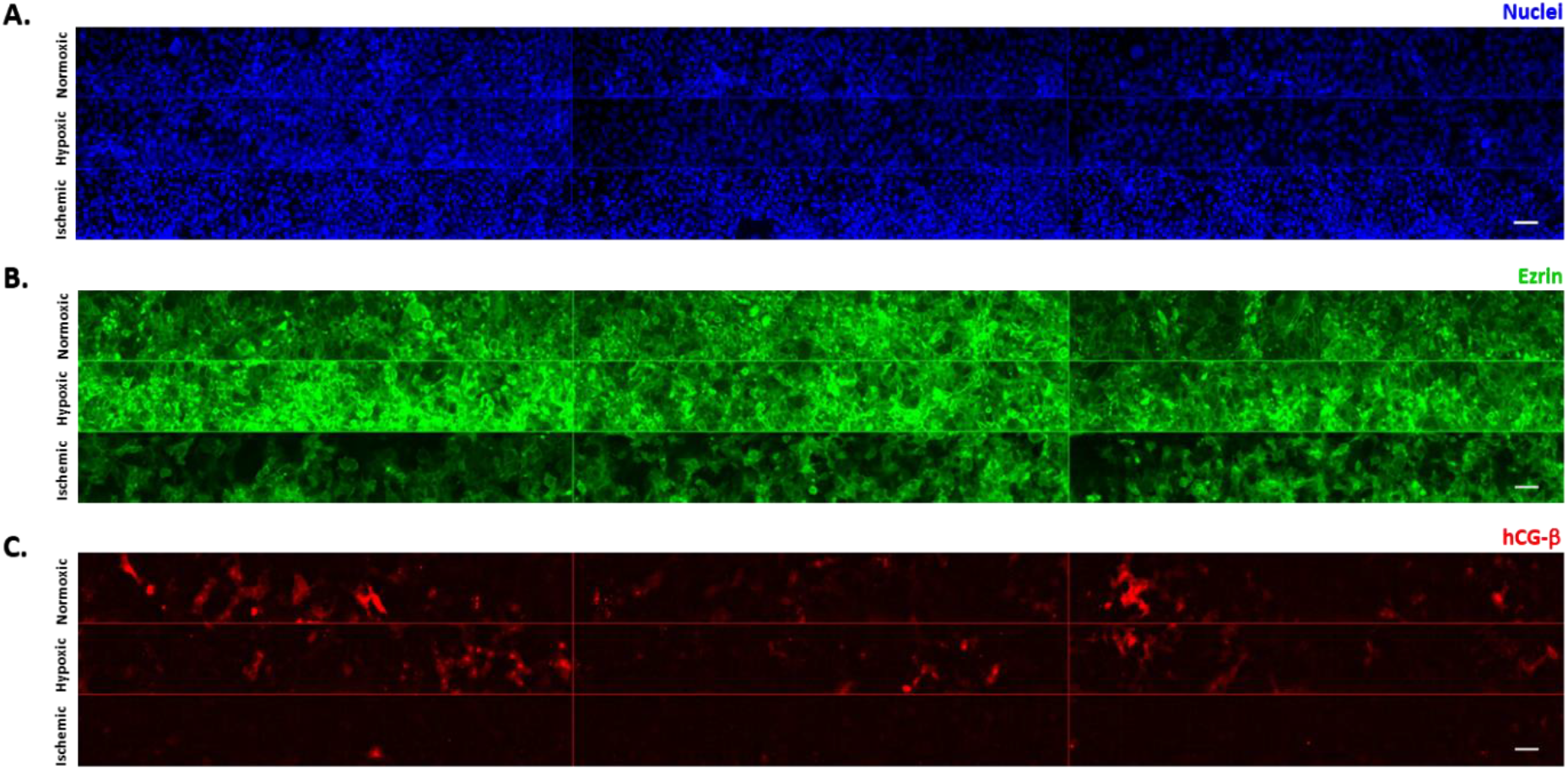
Regions of interest (ROI) for nuclei count and Ezrin, hCG-ß immunostaining quantification. Summary z-projections of **(A)** nuclei stained with Hoechst 33342 (blue), **(B)** Ezrin (green), **(C)** hCG-ß (red) in “Normoxic”, “Hypoxic” and “Ischemic” conditions. Scale bar: 100μm.

**Supplementary figure 5.**
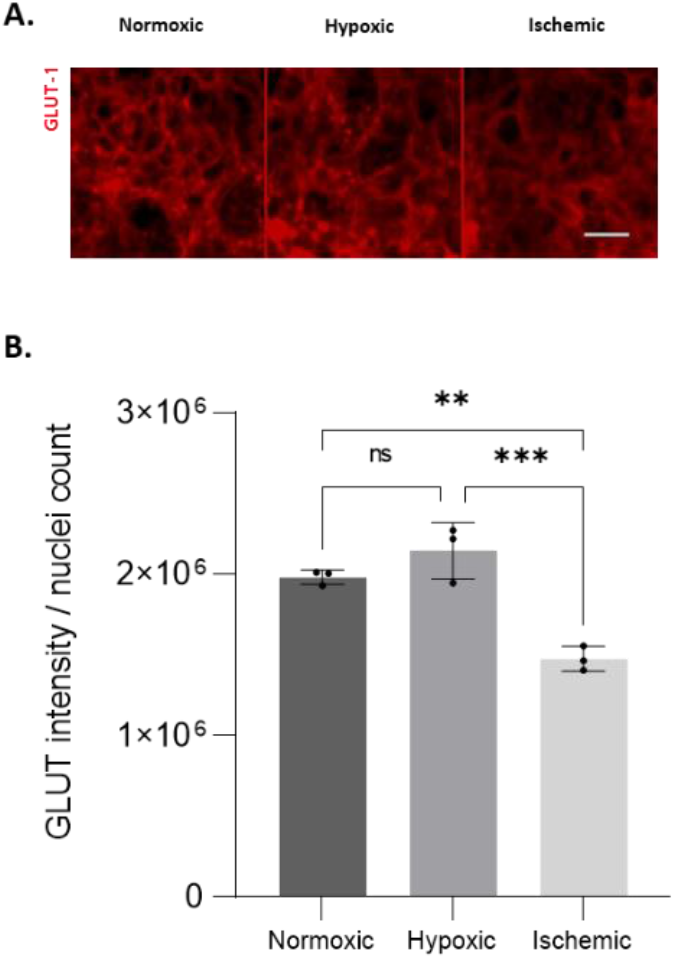
Glucose transporter-1 (GLUT-1) expression in “Normoxic”, “Hypoxic” and “Ischemic” conditions. **(A)** Representative summary z-projections of GLUT-1 intensity in “Normoxic”, “Hypoxic” and “Ischemic” conditions. Scale bar: 50μm. **(B)** Ratio of GLUT-1 intensity over nuclei count in the syncytium, in “Normoxic”, “Hypoxic” and “Ischemic” conditions. Data points represent individual chips. Data are presented as mean ± SD and statistical analysis was performed using ordinary one-way ANOVA. (N=1, n=3). ns: non-significant, **P < 0,01; ***P < 0,001).

